# Too Close to Eat? Solidarity with Animals, Animal Welfare and Meat Consumption

**DOI:** 10.1101/2020.01.30.926592

**Authors:** Albert Boaitey, Michaela Eden, Simon Jette-Nantel

**Affiliations:** Department of Agricultural Economics, University of Wisconsin-River Falls, River Falls, Wisconsin, United States of America

## Abstract

Meat consumption is influenced by a variety of factors including of empathy and feelings on affinity towards farm animals. The goal of this study was to examine the role of solidarity with animals on meat consumption and attitudes towards the treatment of animals. Data was drawn from a sample of 265 respondents in the US. Correlation and mediation analyses were performed. The results of the correlation analysis indicate a moderate but positive correlation between solidarity with animals and proecological beliefs. The association between attitudes towards the treatment of farm animals and antibiotic use and solidarity with animals was also positive. Relative to meat consumption, the results indicate that proecological beliefs and concerns about the treatment of farm animals negatively influenced consumption. The effect of attitudes towards antibiotic use and solidarity with animals on consumption were however fully mediated by proecological beliefs. The results indicate that social identification with animals can play a significant role in food choice. However, its relationship with proecological beliefs implies that holistic approaches are required to address current livestock production practices that are considered unnatural.

## Introduction

Animal welfare appears to be increasingly important in many countries and it even has become an official part of Schwartz’s well-known theory of human values [1]. Concerns about farm animals relate to how animals are treated and kept. These concerns differ across species and can elicit different responses (such as meat avoidance) from consumers. In general, reductions in the consumption of meat are mostly driven by consumers’ personal health and ethical motives [2,3,4]. In relation to animal welfare, consumers may reduce meat consumption to avoid causing harm to farm animals or as a result of concerns about practices considered unnatural [5]. Animal and human welfare attitudes also influence behaviors such as support for animal rights and causes [2,6]. These attitudes are underpinned by different values and motivations including the extent to which consumers feel connected to farm animals [7]. Human-animal relations are multi-faced in nature and aspects such as affinity with animals have been linked with a number of behaviors: attribution of a higher cognitive ability, anthropomorphism and anti-consumption [8, 9, 10]. The pathways through which these attitudes influence consumer behavior in relation livestock production and consumption are however less understood. This paper focuses on a different aspect of human-animal relation i.e., the concept of solidarity with animals. Solidarity with animals is the sense of connectedness to animals as part of the same social group [11, 12]. This psychological bond with farm animals can have significant implications for public attitudes towards different treatment and farm practices, and meat consumption behavior. It is conceivable that solidarity with animals is linked to a broader connection to nature and a proecological orientation. This notwithstanding, not much has been done to examine the links between solidarity with animals and these important aspects of meat production and consumption. This is partly due to the absence of a well-validated metric to measure solidarity with animals. A solidarity with animals’ scale with good psychometric properties has recently been introduced [11]. In this study, we investigate the relationship between solidarity with animals, proecological beliefs, treatment of farm animals and pork consumption using data from a sample of participants in the United States. Our interest in the role of solidarity is informed by evidence from previous studies [6, 2] which suggests that solidarity with animals can influence socially consequential actions.

Amongst the current litany farm animal welfare issues, the use of antibiotics in livestock production presents a unique case because of its ethical and human health impacts. The latter is in relation to the public health risks posed by the veterinary overuse of antimicrobials and the possibility of animal-human transfer of antimicrobial resistance [13, 14]. The Food and Agricultural Organization (FAO) estimates substantial economic and health consequences^1^ resulting from antimicrobial resistance (AMR) if current trends continue [15]. In contrast to the negative human health impacts, the use of antibiotics can enhance animal welfare by reducing or eliminating the pain associated with disease. Inherent in the dual impacts of antibiotic use is the conundrum faced by consumers who may be concerned about the treatment of animals on farms and the possible negative health impacts. It is obvious that feelings of connectedness to animals further complexifies these relationships. Further, we focus on the hog production in the US for two reasons. First, available estimates indicate that antibiotics are significantly overused in pig production in US - 27.1% of all medically sold antibiotics is used in pigs as compared to the 27.6% is used in human medicine [16]. Second, these high levels of antibiotic use are linked to production and husbandry practices such as the high degree of concentration [16]. For the pork industry, insights from this study is particularly relevance given the current changes in the consumption, the need to provide an appropriate level of animal welfare and the emergence of new products such as lab-grown meat often marketed as animal welfare friendly [17].

## Literature review

The subject of the effect of human-animal relationship on consumer behavior has been of long-standing interest because of its influence on a number of attitudes and behaviors [6, 9]. These attitudes can also serve as a motivation for ethical food choice [4]. It has been shown that associating humanlike attributes such as perceived intelligence and appearance to animals influences disgust at the thought of eating meat which leads to meat avoidance [18, 9, 5]. However, the extent to which animals are consider similar humans can dampen or exacerbate these responses [7]. Other studies have looked at the role of psychological commitments to groups such as solidarity with animals, as a determinant of attitudes towards animals [11]. The authors found that solidarity with animals, was negatively associated with meat eating frequency, higher moral concerns for animals, and a greater likelihood to donate to animal charities. This social identity dimension of attitudes towards animals influences other consumer response (such as activism for animal rights) in so-called factory farms which are often considered unnatural [6]. Perceptions about animal welfare are also influenced by both objective and normative judgements [19]. The authors found that animals experiencing negative emotions but living in their natural setting were perceived as experiencing a higher animal welfare standard [19].. This is in comparison to animals in unnatural settings experiencing positive emotions. This may be indicative of linkages between perceived FAW and proecological attitudes and by extension meat consumption. Other studies have examined the effect of proecological attitudes (as measured by the New Ecological Paradigm (NEP) scale) [20], on meat consumption, however, the evidence appear inconclusive [21]. In this study, we explore the relationship between attitudes towards antibiotic use, the treatment of animals, proecological beliefs and solidarity with farm animals. We also model the relationship between these variables and their influence on pork consumption using mediation analysis. The use of mediation models to address similar problems have been reported in the literature [14, 22,]. In this study, the effect of solidarity with animals, attitude towards antibiotic use on meat consumption is assumed to be mediated by proecological beliefs.

## Methods

An online survey instrument designed in Qualtrics was used to collect data used in the study in February and March of 2019. The study adhered to the ethical guidelines and was approved by the Institutional Review Board (IRB) of the University of Wisconsin – River Falls (approval number: H2018 KO10). In total 265 responses were obtained from random of 1,000 adult respondents who provided their email address to the Survey Research Center (SRC) at the University of Wisconsin-River Falls and received an email with a hyperlink to the survey. Out of the total returned, 207 questionnaires were answered completely and were therefore considered usable. Incomplete surveys were considered invalid. The sample included 83% (n=171) females and 17% (n=36) male. The latter is lower than the share of males in Wisconsin which is about 50% (US Census Bureau, 2019). The mean age of participants in this study was 34.4 (SD=16.45), lower than the 38.92 reported in the 2017 US census. The average household income of $68,852 (SD=$3,3283), reported in our survey was however higher than the average income of $56,769 in Wisconsin in 2017 [23]. In general, the differences between the sample and the general population are expected given that most of our respondents were in the River Falls area.

The questionnaire was divided into three subsections. In the first part, respondents were asked to provide information on their pork purchase frequency and their preferences for different pork attributes. The second part of the questionnaire included scale items that measured respondents’ attitude towards farm animals, the use of antibiotics in livestock, the environment etc. This section also included the solidarity with animals scale. The last section of the questionnaire measured respondents’ socio-demographic characteristics. Below is an overview of selected questions included in the present analysis.

To measure the frequency of pork consumption, participants were asked, “How often do you consume pork”. Reponses were rated on an eight-point scale with end points: 0=never, to 7=daily. Three measures of animal welfare attitudes and perceptions were included in the survey: attitude towards the treatment of animals [24] and attitudes towards antibiotic use [25]. A five-point Likert scales with end points, 5=strongly agree to 1= strongly disagree, was used in the animal treatment and antibiotic use scales. Proecological attitudes were measured with a reduced version of the New Ecological Paradigm (NEP) scale, comprising 6 items rated on a 5-point scale (1= strongly disagree; 5=strongly agree) [20]. Three of the six items capture the pro-anthropocentric dominant social paradigm (DSP) whilst the remaining statements capture a proecological orientation (See Table 4).

**Table 1.** Descriptive Statistics and Factor Loadings for Items of Solidarity with Animals Scale.

| Items | This study |  |  | Amiot and Bastian (2017) |  |  |
| --- | --- | --- | --- | --- | --- | --- |
|  | M | SD | Factor loading | M | SD | Factor loading |
| I feel a strong bond toward other animals (Strong bond) | 4.19 | 0.92 | 0.65 | 4.97 | 1.54 | 0.90 |
| I feel solidarity toward animals (Solidarity) | 3.70 | 1.02 | 0.95 | 4.84 | 1.57 | 0.88 |
| I feel close to other animals (Closeness) | 4.03 | 0.95 | 0.80 | 4.57 | 1.83 | 0.93 |
| I feel a strong connection to other animals (Connection) | 4.04 | 0.93 | 0.95 | 4.17 | 1.89 | 0.94 |
| I feel committed toward animals (Commitment) | 4.14 | 0.95 | 0.82 | 4.35 | 1.84 | 0.85 |
| Cronbach's alpha | 0.92 |  |  | 0.94 |  |  |

Note: M=Mean; SD Standard Deviation

**Table 2.** Results of Correlation Analysis: Solidarity with Animals and Attitudes Towards the Treatment of Animals.

|  | It is important to me that animal products I eat have been produced in a way that the animal's rights have been respected | It is important to me that the animal products I eat have been produced in a way that the animals have experienced as little pain as possible | The treatment of animals in livestock farming raises serious ethical questions | Increased regulation of the treatment of animals in farming is needed | In general, humans have too little respect for the quality of life of farm animals | solidarity with animals scale |
| --- | --- | --- | --- | --- | --- | --- |
| It is important to me that animal products I eat have been produced in a way that the animal's rights have been respected | 1 |  |  |  |  |  |
| It is important to me that the animal products I eat have been produced in a way that the animals have experienced as little pain as possible | 0.59** | 1 |  |  |  |  |
| The treatment of animals in livestock farming raises serious ethical questions | 0.15* | 0.18** | 1 |  |  |  |
| Increased regulation of the treatment of animals in farming is needed | 0.23** | 0.26** | 0.75** | 1 |  |  |
| In general, humans have too little respect for the quality of life of farm animals | 0.21** | 0.15* | 0.72** | 0.67** | 1 |  |
| solidarity with animals scale | 0.28** | 0.32** |  | 0.17* | 0.15* | 1 |

Notes: \*\*\*Correlation is significant at the 0.01 level; \*\*Correlation is significant at the 0.05 level;
\*Correlation is significant at the 0.10; insignificant estimates not reported

**Table 3.** Results of Correlation Analysis: Attitudes Towards the Use of Antibiotics and Solidarity with Animals.

|  | The process of developing and testing antibiotics for use in livestock production proves their effectiveness and safety | The use of antibiotics is a better strategy than destroying the affected animals | For serious animal bacterial diseases, requirements for farmers to use antibiotics should be in place | We live in such a hygienic environment that the use of animal antibiotics is redundant | Overall, the use of animal antibiotics delivers more benefits than harm | Use of antibiotics in livestock cannot be seriously harmful; otherwise, authorities would ban the use | There is a good reason why the use of certain animal antibiotics is recommended | solidarity with animals scale |
| --- | --- | --- | --- | --- | --- | --- | --- | --- |
| The process of developing and testing antibiotics for use in livestock production proves their effectiveness and safety | 1 |  |  |  |  |  |  |  |
| The use of antibiotics is a better strategy than destroying the affected animals | 0.60** | 1 |  |  |  |  |  |  |
| For serious animal bacterial diseases, requirements for farmers to use antibiotics should be in place | 0.41** | 0.48** | 1 |  |  |  |  |  |
| We live in such a hygienic environment that the use of animal antibiotics is redundant |  | -0.20** | -0.16* | 1 |  |  |  |  |
| Overall, the use of animal antibiotics delivers more benefits than harm | 0.53** | 0.48** | 0.35** |  | 1 |  |  |  |
| Use of antibiotics in livestock cannot be seriously harmful; otherwise, authorities would ban the use | 0.63** | 0.58** | 0.51** | -0.21** | 0.53** | 1 |  |  |
| There is a good reason why the use of certain animal antibiotics is recommended | 0.60** | 0.49** | 0.44** |  | 0.50** | 0.52** | 1 |  |
| solidarity with animals scale |  |  | 0.17* |  | 0.16* | 0.19** |  | 1 |

**Table 4.** Results of Correlation Analysis: NEP Scale and Solidarity with Animal.

|  | Plants and Animals Have a Right to Exist | Humans Have the Right to Modify Natural Environments | Interference with Nature | Nature Can Cope with Industrial Nations | Humans Abuse Environment | Ecological Crisis is Exaggerated | solidarity with animals scale |
| --- | --- | --- | --- | --- | --- | --- | --- |
| Plants and Animals Have a Right to Exist | 1 |  |  |  |  |  |  |
| Humans Have the Right to Modify Natural Environments | -0.20** | 1 |  |  |  |  |  |
| Interference with Nature | 0.33** | -0.30** | 1 |  |  |  |  |
| Nature Can Cope with Industrial Nations | -0.19** | 0.35** | -0.26** | 1 |  |  |  |
| Humans Abuse Environment | 0.31** | -0.26** | 0.47** | -0.39** | 1 |  |  |
| Ecological Crisis is Exaggerated | -0.34** | 0.17* | -0.26** | 0.48** | -0.53** | 1 |  |
| solidarity with animals scale | 0.27** |  |  |  |  | -0.17* | 1 |

Notes: \*\*\*Correlation is significant at the 0.01 level; \*\*Correlation is significant at the 0.05 level;
\*Correlation is significant at the 0.10; insignificant estimates not reported

Following [11], a five-item scale was used to measure solidarity with animals. The scale comprised of the following items each rated on a seven-point scale with endpoints, 1=strongly agree;7=strongly disagree: “I feel a strong bond toward other animals”; “I feel solidarity toward animals”; “I feel close to other animals”; “I feel a strong connection to other animals”; and, “I feel committed toward animals”.

### Statistical Analyses

Multiple approaches were used to analyze the data. These include principal components analysis (PCA), independent sample t-tests and mediation analysis. The PCA approach was used to extract factors form the multi-items. Pearson correlation coefficient between the solidarity with animals scale and the: attitudes towards the treatment animals, antibiotics use in livestock statements and and proecological were also estimated. The PCA and correlation analysis were performed in performed using IBM SPSS software (version 24). Mediation analyses were conducted to assess the total effect of solidarity with animals on meat consumption frequency in addition to the extent to which this effect is mediated by proecological beliefs. We also estimated the mediated moderation pathway (indirect effect of attitudes towards antibiotics use and the treatment of animals) on consumption via proecological beliefs.

## Results

The analysis is preceded by a descriptive analysis of the scales included in the study. From Table 1, the mean scores (M), factor loadings and standard deviations of scale items of the solidarity with animals scale were comparable to the original application of the scale. The estimates in this study were however marginally lower. The alpha coefficients were high (>0.90) in both cases suggesting that the items have a high internal consistency. Consistent with the findings of [11], the mean scores of female participants (4.10) was significantly higher (t=2.36;P-value 0.02) than those of male participants (3.68). This supports the identified gender differences in solidarity with animals reported in [11].

Most respondents (53%) strongly agreed with the statement that the use of antibiotics is a better strategy than destroying the affected animals (Fig 1). In contrast, most respondents (23%) were in strong disagreement with that statement that, overall, the use of animal antibiotics delivers more benefit than harm. Respondents were somewhat adverse (16% strongly disagree) to regulated use of antibiotics for serious bacterial disease. However, they seem to recognize the need for antibiotics use in livestock and believed that the process of development and testing of antibiotics was generally effective and safe (77% agree and strongly agree).

**Fig 1.**
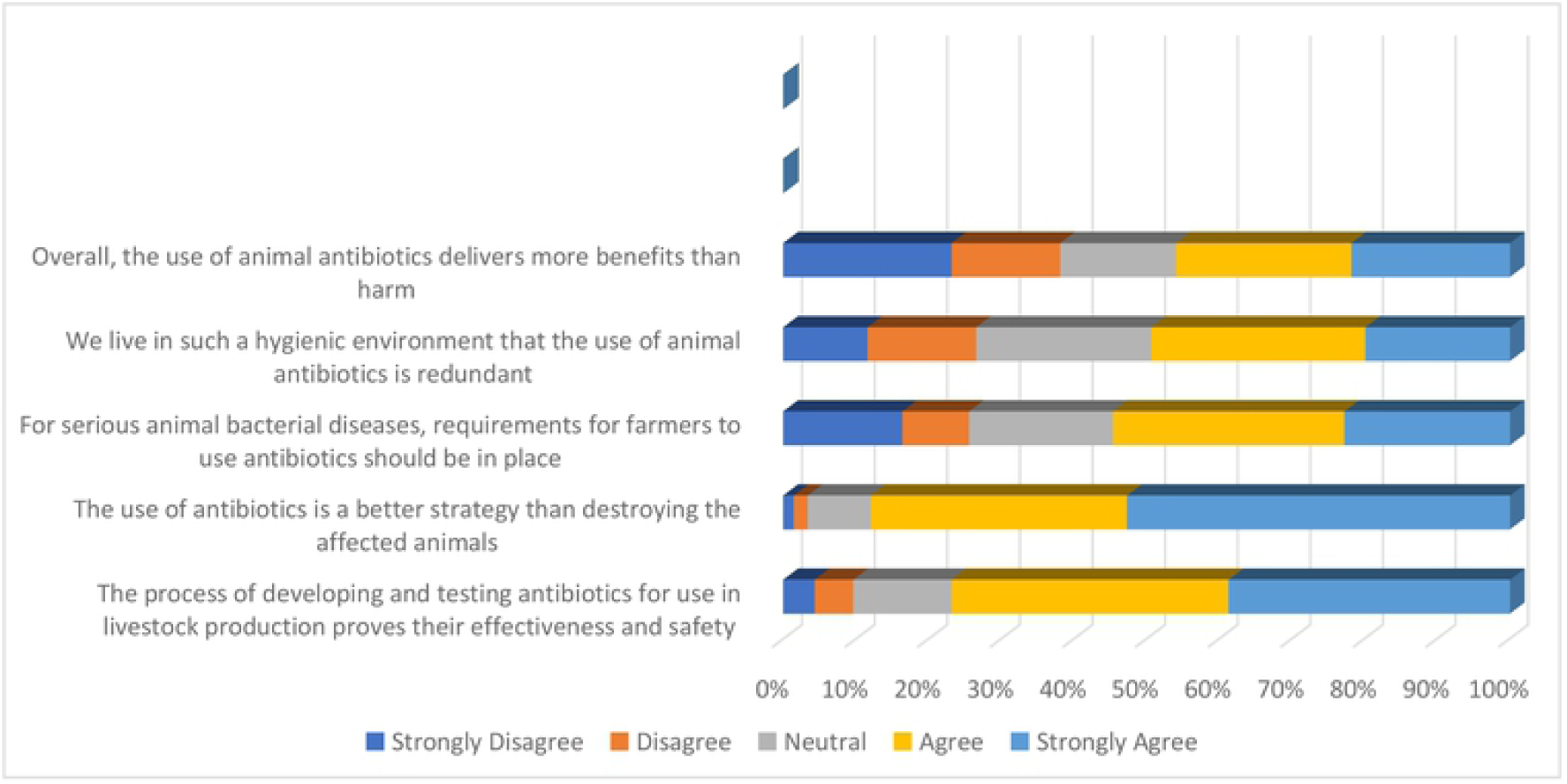
Participants attitudes towards the use of antibiotics in livestock.

Fig 2 shows the participants agreement with various attitudes towards the treatment of animals statements. The first three statements relate to ethical aspects of the treatment of animals whilst the remaining two assesses the role of animal welfare in food choice [24]. In general, respondents expressed higher levels of agreement with the latter statements as compared to the former three. Most respondents strongly agreed (53%) and agreed (35%) that the animal products they consumed are produced by animals that have experienced as little paid as possible. Another 77% strongly agreed (39%) and agreed (38%) with the statement that it was important to them that the products they consumed were sourced from animals whose rights have been respected. In contrast, a lower proportion of respondents agreed and strongly agreed with statements: humans have little respect for animals (46%); increased regulation of the treatment of animals is needed (49%); and, the treatment of animals raises serious ethical questions (55%). These trends are generally consistent with a estimates in Canada although the magnitudes differ (for example see [26]).

**Fig 2.**
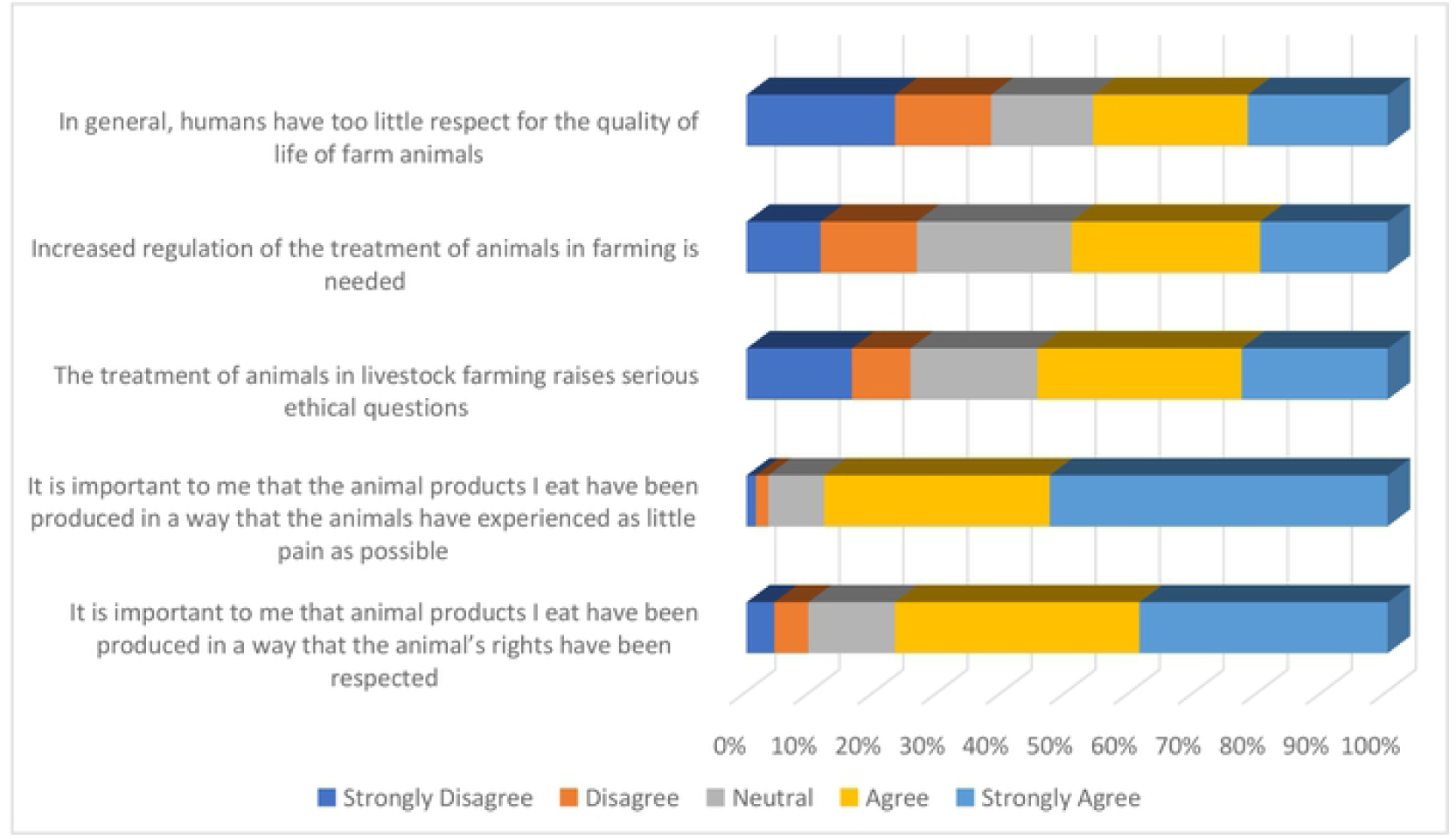
Participants attitudes towards the treatment of farm animals.

### Correlation analysis

The Pearson correlation analysis to assess the relationship between the solidarity with animals scale and attitudes towards the treatment of animals and the use of antibiotics are reported in Tables 2 and 3 respectively. The results indicate that there is positive significant correlation between the solidarity with animals and various items in the animal attitudes scale (Table 2). The correlations are however moderate to weak depending on the item. The stronger correlation was between the ethical treatment of animals statements, “It is important to me that animal products I eat have been produced in a way that the animal’s rights have been respected “(r=0.28); and, “It is important to me that the animal products I eat have been produced in a way that the animals have experienced as little pain as possible” (r=0.32). In contrast, correlations when positive was weaker for statements that assess perceptions towards animal welfare in food choice: increased regulation of the treatment of animals in farming is needed”(r=0.17); and, “In general, humans have too little respect for the quality of life of farm animals” (r=0. 15). These results suggest a positive relationship between solidarity with the animals and perceptions about farm husbandry practices. The relationship is stronger for the ethical treatment issues as compare to the role of animal welfare in food choice.

From Table 3, relationship between solidarity with animals and attitudes towards the use of antibiotics shows marked differences across the various items of the latter scale. Out of the seven items of the antibiotics use scale, two items, i.e. “We live in such a hygienic environment that the use of animal antibiotics is redundant” and “There is a good reason why the use of certain animal antibiotics is recommended”, were not significantly correlated solidarity with animals scale. statements on the trust in in regulation of antibiotic use, relative net benefit of antibiotic use and need for the mandated use of antibiotics to treat serious disease were all positively correlated with the attitudes towards the use of antibiotics. Overall, these results suggest a positive but weak association between solidarity with animals and support for the use of antibiotics in livestock

We also assessed the correlation between the solidarity with animals and environmental beliefs (Table 4). The results indicate that solidarity with animals is positively associated with selected proecological beliefs and negatively correlated with anthropocentric beliefs. This is evident from the positive correlation between the solidarity scale the proecological statement, “Animals and plants have a right to exist”, and the negative correlation with the anthropocentric statement, “The ecological crises is exaggerated”. The association was stronger (r=+2.27) in the case of the former than the latter (r=−0.17).

### Mediation Analysis

The mediation analysis is preceded by an overview of self-reported pork consumption behavior amongst survey participants. Most participants (37%) reported consuming pork 3-4 times a week. A smaller proportion (3%) of participants reported eating pork daily or never (4%). Table 5 is a summary of the results of the mediation model estimated.

**Table 5:** Results of Mediation Analysis.

| **Outcome [Consumption]** | *Direct effect (DE)* |  | *Total effect (TE)* |  |
| --- | --- | --- | --- | --- |
|  | Coefficient | SE | Coefficient | SE |
| NEP | -0.27*** | 0.10 | -0.27*** | 0.10 |
| solidarity with animals | 0.04 | 0.10 | 0.01 | 0.10 |
| attitudes towards antibiotics use | 0.02 | 0.10 | 0.06 | 0.10 |
| attitude towards the treatment of animals | -0.23** | 0.11 | -0.36*** | 0.10 |
| **Outcome [NEP]** |  |  |  |  |
|  | *Direct effect (TE)* |  | *Total effect (TE)* |  |
|  | Coefficient | SE | Coefficient | SE |
| solidarity with animals | 0.11* | 0.06 | 0.11* | 0.06 |
| attitudes towards antibiotics use | -0.13** | 0.06 | -0.13** | 0.06 |
| attitude towards the treatment of animals | 0.46*** | 0.07 | 0.46*** | 0.67 |

Note: \*\*\*, \*\*, \* denote significance at 1%; 5% and 10% respectively; SE is standard error

Attitudes towards the treatment of animals and proecological beliefs. (NEP) have a negative effect on consumption. The direct effect and total effect of the former were −0.23 and −0.36 respectively. This suggests that part of the effect of attitudes towards the treatment of animals on consumption is mediated by proecological beliefs. Solidarity with animals did not have a direct effect on consumption. However, its effect on proecological beliefs was positive (0.11). This suggests that effect of solidarity with animals on pork consumption is fully mediated by proecological beliefs. Attitudes towards antibiotic use in livestock had no direct effect on meat consumption. It was negatively associated (−0.13) with proecological beliefs. Considering that the use of antibiotics may be considered unnatural, this result is unsurprising.

In sum, the mediation analyses suggest attitudes concerns about livestock husbandry practices and proeclogical beliefs reduce the tendency to consume meat. The effect of solidarity and attitudes towards antibiotic on meat consumption is fully mediated by proecological beliefs.

## Discussion

In this study we set out to investigate relationship between solidarity with animals, meat consumption and attitudes towards production practices. Specifically, we focused on attitudes towards the treatment of animals and the use of antibiotics. We found a positive but moderate association between solidarity with animals and these attitudes. This suggests individual with a stronger sense of social identification with farm animals may be more receptive to the use of antibiotics in livestock and proper care of farm animals. Solidarity with animals was also positively associated with proecological beliefs. The results further suggest the effect of solidarity with animals and attitudes towards antibiotic use on consumption is mediated by proeological beliefs. The indirect effect of solidarity with animals on consumption corroborates evidence from previous studies on the impact of human-animal on meat consumption [5, 22,28] The additional insights provided in this study is that, the effect of feelings of connection to animals is perhaps subsumed under a broader feeling of connectedness to nature. This outcome seems consistent with the relationship between solidarity with animals and speciesism reported in [11] and the role of animal welfare and ecological concerns in food choice [6, 27]. An important implication is that while solidarity with animals may invoke a higher degree of acceptance for practices that ameliorate the pain of farm animals, meat consumption may reduce their consumption of meat if these practices (e.g. the use of antibiotics) are considered unnatural. Given that the respondents in our sample are mostly non-vegetarian, the effect of proecological beliefs and attitudes towards the treatment of animals on consumption is indicative of possible conscientious omnivore behavior [29]. Where, respondents may not completely shift away from meat production but may purchase meat from ethical sources. For the conventional pork industry, potential negative impacts of these attitudes on consumption can be attenuated by addressing the concerns about antibiotics use as part of a broader range of ethical considerations (environmental and animal welfare). Partial approaches may be less successful given the higher order linkages identified in the present study. Our results should be interpreted with a few caveats in mind. Our sample overrepresents female respondents (80%) and this can be a source of bias. We also did not consider other meat consumption behaviors-vegetarians, flexitarian, omnivores It is plausible that the role of solidarity with animals across the different consumption behavior and other livestock species [10, 19] may be different. These limitations are avenues for future research.

